# The neural representational space of social memory

**DOI:** 10.1101/130351

**Authors:** Sarah L. Dziura, James C. Thompson

## Abstract

Social functioning involves learning about the social networks in which we live and interact; knowing not just our friends, but also who is friends with our friends. Here we utilized a novel incidental learning paradigm and representational similarity analysis (RSA), a functional MRI multivariate pattern analysis technique, to examine the relationship between learning social networks and the brain's response to the faces within the networks. We found that accuracy of learning face pair relationships through observation is correlated with neural similarity patterns to those pairs in the left temporoparietal junction (TPJ), the left fusiform gyrus, and the subcallosal ventromedial prefrontal cortex (vmPFC), all areas previously implicated in social cognition. This model was also significant in portions of the cerebellum and thalamus. These results show that the similarity of neural patterns represent how accurately we understand the closeness of any two faces within a network, regardless of their true relationship. Our findings indicate that these areas of the brain not only process knowledge and understanding of others, but also support learning relations between individuals in groups.

**Significance Statement:** Knowledge of the relationships between people is an important skill that helps us interact in a highly social world. While much is known about how the human brain represents the identity, goals, and intentions of others, less is known about how we represent knowledge about social relationships between others. In this study, we used functional neuroimaging to demonstrate that patterns in human brain activity represent memory for recently learned social connections.

---

Social relationships guide and support much of human behavior. Not only do we form strong lifelong family bonds, we also interact with others in work, education, and leisure settings and create lasting non-kin relationships. For many species, including humans, non-kin based social networks can have important consequences for health and fitness (Cheney, 2011; Tung et al., 2015). Benefits of social relationships can come both from direct connections (our friends) as well as second-order or indirect connections (friends of our friends) (Brent, 2015; Seyfarth & Cheney, 2015). A considerable amount of research has revealed the cognitive and neural mechanisms underlying the representation of social faces, voices, and bodies (Allison, Puce, & McCarthy, 2000; Arsenault & Buchsbaum, 2015). There is also a good understanding of the neural basis of social knowledge about other individuals (Koski, Xie, & Olson, 2015; Wang et al., 2017), how we represent ingroup versus outgroup members (Van Bavel & Cunningham, 2012; Shkurko, 2013), and how we represent the mental states of others (Contreras, Schirmer, Banaji, & Mitchell, 2013; Saxe & Kanwishwer, 2003).

Less well understood is how we represent knowledge about indirect social connections. Memory for who knows whom is important for navigating social relationships, including knowing who to approach for information, support, and other resources. Individual differences in sociality have been linked to neural and behavioral measures of social perception (Baron-Cohen et al., 1999; Dziura & Thompson, 2014; Kanai, Bahrami, Roylance, & Rees, 2012), and there are differences in how well people can remember social networks (Brewer, 2000; Casciaro, 1998; Freeman, Romney, & Freeman, 1987). However, the underlying neural processes involved in learning complex social network relationships has not been thoroughly examined. Human social organization is dynamic, as the composition of groups and the ties within them change over an individual's lifetime (Couzin, 2006). Prior literature indicates several brain areas likely to be important for representing information about social networks. Perception of changes in relationship ties and tie strength has been linked to activity in the bilateral posterior superior temporal sulcus (pSTS) and temporoparietal junction (TPJ) (Bault, Pelloux, Fahrenfort, Ridderinkhof, & Van Winden, 2015). Retrieval of social knowledge, including consideration of kin group cohesion, involves medial and lateral prefrontal cortex (mPFC; lPFC) (Rüsch et al., 2014; Satpute, Badre, & Ochsner, 2014). Learning and representing information about social hierarchy, an important component to many social networks, recruits amygdala, hippocampus, and ventral mPFC (Kumaran, Melo, & Duzel, 2012). A recent study by Parkinson and colleagues (2017) revealed that the similarity of local patterns of fMRI responses in ventral mPFC and lPFC, as well as lateral temporal cortex and TPJ, to viewing videos of individuals from participants’ real world social network, conveyed information about network position of the members. These findings suggest that information about social network relationships is represented in patterns of fMRI responses associated with viewing individuals from one's network.

In this study, we examined the memory and neural representation of connections between members of two novel social networks, using fMRI and representational similarity analysis (RSA). Artificial networks were used in order to experimentally control the closeness of network members and assess the role of the memory for relationship strength in fMRI responses. We examined if the pattern similarity of fMRI responses to any two faces from a learned social network reflected the tie strength (closeness) of those two individuals within the network: that is, does the similarity of the pattern of response to two network members increase as a function of the closeness of those members? We also examined if the memory for tie strength between network members was related to the similarity of the fMRI voxel pattern response to the faces of members. To understand the contribution of the frequency of face pairing during network learning to memory and neural representations, we compared a network in which centrality differed between members (i.e. some members had more connections than others) to a network with no individual centrality.

## Materials and Methods

### Participants

22 healthy individuals (10 females; age range = 18-34; mean age = 23; ethnicity = 64% White, 18% Hispanic/Latino, 18% Asian) participated in a 1.5 hour learning session immediately followed by a 1.5 hour fMRI scanning session. Behavioral data from a total of 31 individuals was collected, but seven subjects did not meet the learning criteria from the behavioral task, one subject was unable to be scanned, and one subject's fMRI data was incomplete. All participants were right handed (self-reported) with normal or corrected-to-normal vision. Participants provided written informed consent in accordance with the Declaration of Helsinki and the Human Subjects Review Board at George Mason University and were compensated for their time.

### Experimental Design and Statistical Analysis Stimuli

Task stimuli consisted of 24 faces of varying ethnicities, equally divided by gender. Faces were all in color and facial expressions were all smiling. These stimuli were downloaded from the Park Aging Mind Laboratory Database at UT Dallas (Minear & Park, 2004) and were chosen to be as realistic to a college campus as possible, ensuring the perception of real people who might interact and be friends with each other.

### Task Design

Participants completed a two-alternative forced choice task to become familiar with the structure of two six-person social networks (Figure 1). Pairs of faces represented connections within each network, with the frequency of pairing indicating relationship strength. Each network had an equal number of male and female faces of varying ethnicities. Network properties differed between the two in that although each network had an equal number of connections of each strength level, there were differences among the individual members (faces) in each network. The faces in network 1 had varying numbers of connections and therefore each had a different average closeness to the rest of the network, whereas the faces in network 2 had an equal number of connections and an equal average closeness to all other faces in the network. This meant that in network 1 the centrality of members was varied (variable-centrality network), while in network 2 centrality was equated across members (fixed-centrality network). This also meant that the frequency of presentation of each face differed in network 1, but was equivalent in network 2. Each trial consisted of a face pair presented for 4 seconds accompanied by a question, and participants were asked to make a comparison between the faces and decide which person better fit the question. Questions consisted of behavioral and personality characteristics taken from various personality surveys included in the International Personality Item Pool (http://ipip.ori.org/). Half of the questions asked which person was more likely to exhibit a characteristic, and half asked which person was less likely (example: “Who is more likely to be easily intimidated?”). Network learning took place in alternating blocks, where the subjects viewed 36 randomly presented trials of one network followed by 36 trials of the second network. Participants completed 720 trials in total (360 per network), with the weakest network connections being presented a total of 20 times and the strongest a total of 80 times.

**Figure 1.**
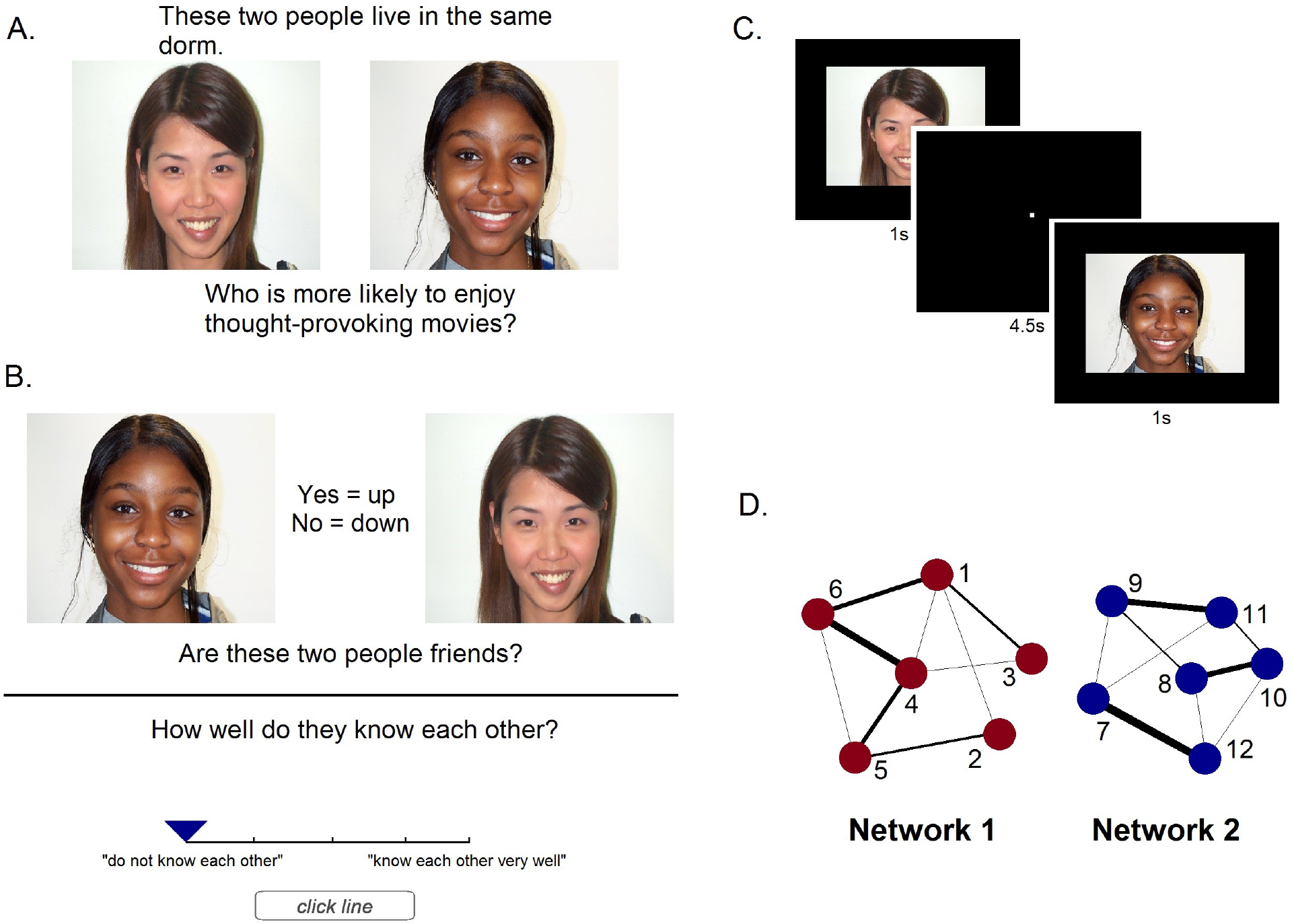
A Example trial of the paired presentation of a social network tie, where subjects were asked to judge between the two faces on an unrelated characteristic. B. Example trial of the recall task, where subjects were asked to report whether a pair of faces was connected, and how well they know each other (0-4 scale). C. fMRI task, where each face was presented individually for 1 second (4.5 second inter-stimulus interval). D. Structure of the two social networks. Each node represents a different face and line thickness represents connection strength. All ties are non-directed (reciprocal).

After completing the paired face viewing portion, participants were explicitly tested on their knowledge of the network connections. They were told that the faces represented college students living in a dorm together, the faces that they saw paired together previously represented friend connections, and the more often they were presented together, the closer in friendship the pair was. They were asked to group all of the faces into two separate halls, as a check to make sure that they could distinguish between the faces in different networks. They were then presented with all possible within-network face pairs twice and asked to rate their relationship on a scale of 0 (do not know each other) to 4 (know each other very well). They were not asked about cross-network face relationships. This explicit testing period was included to ensure that participants learned the structure of the networks to an appropriate level before being scanned. Participants who were within 2 standard deviations of pilot data (hit rate = 0.85, SD = 0.14; false alarm rate = 0.35, SD = 0.15) were included in further analysis. Both parts of the behavioral task (learning and recall) were presented to the participant using PsychoPy version 1.842 software (http://www.psychopy.org/).

The fMRI task stimuli included the same 12 faces from the behavioral task as well as 12 novel faces as a control. Faces were presented one at a time for one second on a black background with a 4.5 second inter-stimulus interval (black screen with a white fixation dot), and participants completed a 1-back task to ensure they were attentive. The task consisted of four runs of 9.6 minutes each, resulting in each face being presented a total of 16 times (not counting repeats, which were included in analysis as a separate regressor). Following the face task, participants underwent an unrelated dynamic localizer session. Localizer stimuli consisted of 18 second blocks each of faces, body parts, outdoor scenes, moving objects, and scrambled objects. The fMRI experiment was presented to the participant using Neurobehavioral Systems Presentation version 16.3 (https://www.neurobs.com).

### fMRI data acquisition, preprocessing, and analysis

fMRI data were collected with a Siemens Allegra 3T scanner and a quadrature birdcage head coil at the Department of Psychology at George Mason University. Visual stimuli were displayed on a rear projection screen and viewed by participants on a head coil-mounted mirror. Blood oxygenation level dependent (BOLD) data were acquired using gradient-echo, echoplanar imaging scans: 40 axial slices (3-mm slice thickness), repetition time (TR)/echo time (TE) = 2350/30 ms, flip angle = 70, 64 x 64 matrix, field of view = 192 mm. 245 volumes were collected in each run. At the end of the fMRI scanning session, one T1 whole-head anatomical structural scan was acquired using a three-dimensional, magnetization-prepared, rapid-acquisition gradient echo (MPRAGE) pulse sequence. The following parameters were used for these scans: 160 1-mm slices, 256 x 256 matrix, field of view = 260 mm, TR/TE = 2300/3.37 ms. Functional data were analyzed using FSL (version 5.0.8) fMRI Expert Analysis Tool (fsl.fmrib.ox.ac.uk) and Matlab (version R2012a) software (http://www.mathworks.com). Preprocessing included brain extraction, high-pass filtering at 96 s, slice-timing correction, motion correction, and smoothing with a 6 mm FWHM kernel. Runs with > 1 mm of motion were run through the BrainWavelet Despiking program in Matlab (Patel et al., 2014). For first-level analysis, linear regression was conducted at each voxel, using generalized least squares with a voxel-wise, temporally and spatially regularized autocorrelation model, drift fit with Gaussian-weighted running line smoother. For second-level analysis, linear regression was conducted at each voxel, using ordinary least squares.

### Regions of Interest (ROI) and Mask Creation

Localizer data preprocessing steps were identical except the functional data was registered only to each subject's specific structural image. Face-selective regions of interest (ROIs) were created from subtracting the combined object, scrambled object, and scene conditions from the face condition. These regions included bilateral posterior STS and fusiform face area (FFA). Activity was thresholded at Z > 3.7 (p < 0.0001) for most ROIs, although this threshold was relaxed to Z > 3 (p < 0.001) in one subject, Z > 2.3 (p < 0.01) in four subjects, and Z > 1.65 (p < 0.05) in three subjects because of lower overall BOLD activity. These masks were projected back into native functional space for further analysis. Finally, an anatomical mask of areas involved in memory for faces (encompassing the bilateral pSTS, extrastriate body area (EBA), ventral temporal/fusiform gyrus, precuneus/posterior cingulate cortex (PC/PCC), and hippocampus) was created from the Harvard-Oxford Cortical Structural Atlas in FSL.

### Univariate Analysis

Each subject's functional data was registered to his or her anatomical scan and then registered to the MNI standard template. The regressors used in the generalized linear modeling (GLM) analysis were Network 1 v. rest, Network 2 v. rest, Control v. rest, and Response Trials v. rest. Contrasts used were Network 1 v. Control, Network 2 v. Control, Both Networks v. Control, Control v. Both Networks, Network 1 v. Network 2, and Network 2 v. Network 1. Group nonparametric 1-sample (conditions v. rest) and 2-sample (condition A v. condition B) t-tests (5000 permutations) including threshold-free cluster enhancement and variance smoothing of 8 mm were conducted with fslrandomise within the mask created from anatomically-defined regions selective for face processing and memory.

### Representational Similarity Analysis

Representational similarity analysis (RSA) is a form of multivariate pattern analysis that compares the distance between stimuli in neural representational space (Kriegeskorte, Mur, & Bandettini, 2008), and correlates this neural information with external patterns of information. In this way it can be utilized to assess different models or patterns of cognition above and beyond univariate analysis, or even more traditional multivariate pattern classification techniques (Haxby, Connolly, & Guntupalli, 2014). For initial analysis of task data, no registration to structural or functional data was carried out, and the smoothing kernel used was 4 mm FWHM. All other preprocessing parameters mirrored the univariate whole-brain analysis. The GLM included separate regressors for each of the 24 faces and repeats. Resulting z-statistics were grouped by network for further analysis. Four separate dissimilarity matrices (DMs) were created for each network (for examples, see Figure 5): true network structure (created from tie strength), perception of network structure (taken from each subject's behavioral recall data after learning the networks), group average of perceived structure (where each face pair's perceived strength was averaged across subjects), and recall accuracy (measured by calculating the absolute distance between the true strength of each face pair and the average strength of the pair reported in the recall phase). The CoSMoMVPA toolbox in Matlab was used for RSA calculations (Oosterhof, Connolly, & Haxby, 2016).

Separate whole-brain searchlights using Spearman correlations (size = 50 voxels) were conducted on the average z-statistics for the faces within each network for each DM. The ensuing correlation maps were transformed into standard space for group analysis. No significant group differences were found across the two networks (in group nonparametric paired-sample t-tests with 5000 permutations), so the correlation maps in individual subject space were then averaged across networks within subjects and transformed again to standard space for across-network group analysis. Group nonparametric 1-sample t-tests (5000 permutations) including threshold-free cluster enhancement and variance smoothing of 8 mm were conducted with fslrandomise. Resulting t-statistic maps were visualized in the MNI volume as well as transformed to the PALS-B12 standard atlas in Caret (http://www.nitrc.org/projects/caret/) for surface data visualization (Van Essen, 2005). RSA was also carried out within each localizer-defined ROI and the resulting correlations within each region were averaged across subjects.

## Results

### Behavioral task

Participants became familiar with the structure of two six-person social networks by viewing two faces presented simultaneously (See Figure 1). A paired set of faces represented a connection within the network, with the frequency of pairing indicating relationship strength. Analysis of social network recall data was conducted in Microsoft Excel (version 2016) and R Version 3.3.2 (https://www.r-project.org/). Subjects correctly identified relationship ties significantly greater than chance across both networks (t(21) = 8.08, p = 7.004e-08). Table 1 shows the average hit rate, false alarm rate, sensitivity (d′), and the correlation between true and reported perceived strength for ties and relationship strength across subjects. Paired sample two-tailed t-tests revealed no significant differences between recall measures for the two networks. There were also no significant age or gender effects for any of the measures. When averaged together across subjects, group perceived relationship strength was highly correlated with the true network structure (r = 0.896, p < 0.00001). In order to assess whether our behavioral task was comparable to previous forms of social network learning and recall, we calculated performance measures used by Brashears (2013). Accuracy refers to the number of ties correctly recalled divided by the number of total ties reported, coverage refers to the number of ties correctly recalled divided by the total tie number in the network, and performance refers to the product of accuracy and coverage. T-tests revealed no significant differences between accuracy or performance measures in our task and those of Brashears (accuracy: t(21) = 0.98, p = 0.34; performance: t(21) = 0.58, p = 0.56), and we actually saw an increase in coverage (t(21) = 3.58, p = 0.002), although our networks were smaller, so participants did not need to remember as many ties.

**Table 1.** Accuracy of recalling network relationships after incidental learning. Hit rate, false alarm rate, and d’ represent the accuracy of recalling the true connections within the networks. Strength correlation refers to the correlation between the matrix of true relationship strength of the faces in the networks and the behavioral judgments of strength, and is therefore a measure of accuracy of recalling relationship strength. T-values and p-values for paired sample two-tailed t-tests between the two networks are reported at the right of the table. Bold indicates primary data, and italics indicate the standard deviation of the data.

|  | **Network 1** | **Network 2** | **Both Networks** | **t-value (p)** |
| --- | --- | --- | --- | --- |
| **Hit Rate** | **0.82** | **0.83** | **0.83** | **-0.13 (0.89)** |
| *SD* | *0.14* | *0.11* | *0.09* |  |
| **False Alarm Rate** | **0.39** | **0.47** | **0.43** | **-1.23 (0.23)** |
| *SD* | *0.21* | *0.24* | *0.19* |  |
| **d'** | **1.3** | **1.2** | **1.2** | **0.70 (0.49)** |
| *SD* | *0.71* | *0.86* | *0.6* |  |
| **Strength Correlation (r)** | **0.58** | **0.53** | **0.54** | **0.82 (0.42)** |
| *SD* | *0.25* | *0.21* | *0.21* |  |

When exploring network recall, it is important to not only look at the correctly identified ties, but also at the pattern of mistakes made. Specifically, we wanted to see whether there are systematic biases that could be predicted by the level of relationship strength of the friend pairs. We assessed recall by relationship strength by looking at the relative direction of the errors made (i.e. how much subjects overestimated or underestimated the strength of the connection). A linear mixed effects regression model (fixed effect = strength; random effects = subject, residual) revealed that relationship strength affected recall error compared to a null model (χ^2^(1) = 226.9, p < 2.2e-16). This pattern shows that overall, weak ties were reported to be stronger than they actually were whereas strong ties were reported to be less strong (Figure 2a). This reflects a general tendency to assume a mid-level relationship between observed people when the relationship is not explicitly known or is unable to be recalled. This central tendency effect seems to be robust, as it was also observed in a separate subject sample (N = 23, 17 females, mean age = 19.6 (sd = 2.4)) learning a larger social network (N= 9) and a larger possible range of relationship tie strengths to choose from (0-6) (χ^2^(1) = 362.84, p < 2.2e-16) (Figure 2b). In order to be able to compare network memory performance to the neural patterns in response to each face in the network, we converted the relative error for each subject to absolute error, which gives a measure of distance from the true network structure, regardless of the direction of that error. The absolute error measure for each subject for each network was then used as a dissimilarity model for RSA to elucidate what neural patterns underlie these errors.

**Figure 2.**
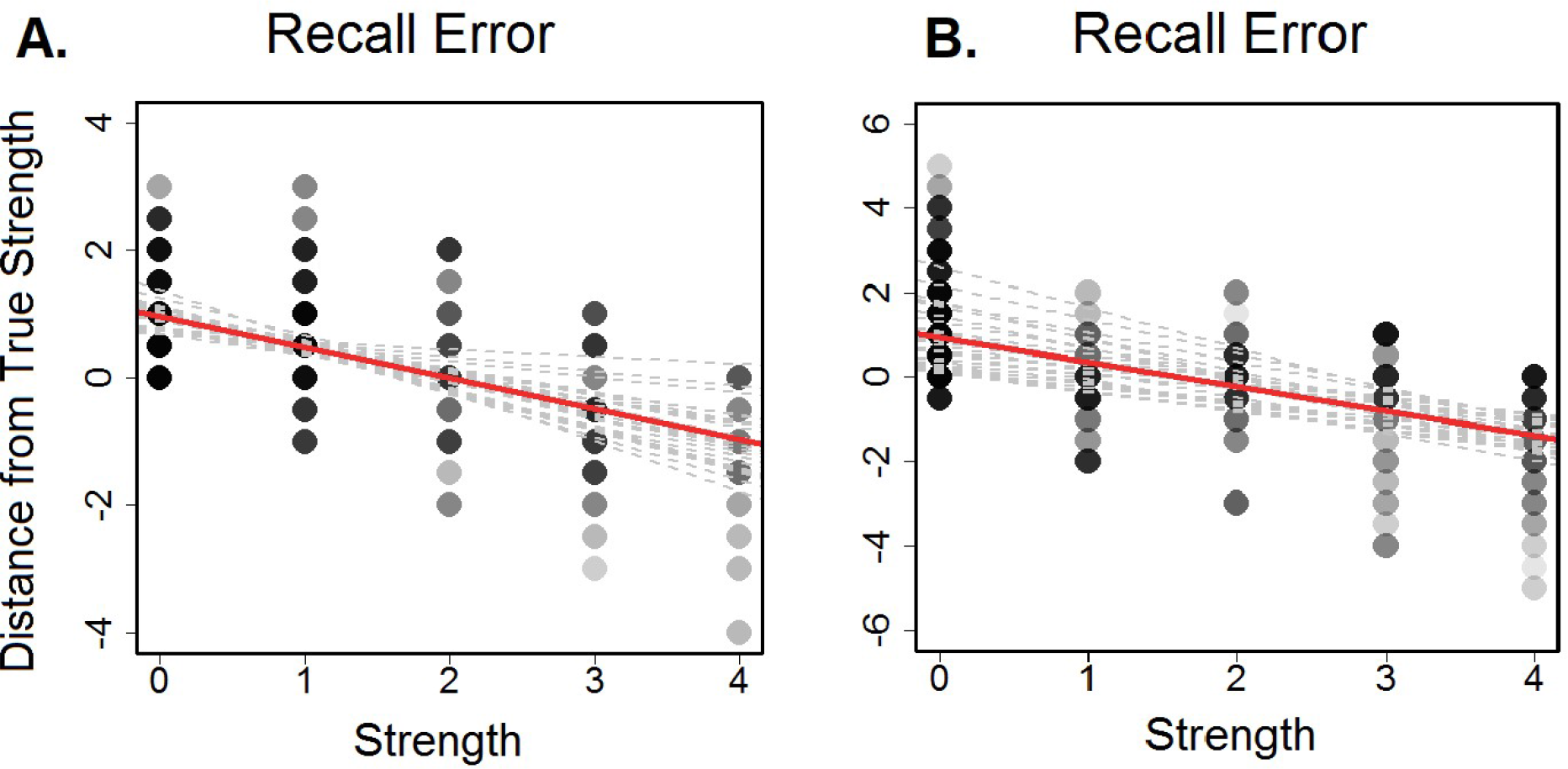
A Each subject's error by pair strength level (0 = unconnected, 4 = close friends) from the primary dataset. Positive values = overestimation of strength and negative values = underestimation of strength. Gray lines show individual subject regression lines. The red line shows the group regression line. B. Subject error by pair strength level (0 = unconnected, 6 = close friends) from the secondary dataset with a larger social network (N = 9). Positive values = overestimation of strength and negative values = underestimation of strength. Gray lines show individual subject regression lines. The red line shows the group regression line.

### fMRI Results

During fMRI scanning, participants viewed the original faces from the social network behavioral session, as well as 12 novel faces and were asked to press a button when they saw a face repeated to guarantee attention. We first conducted a GLM comparing the 12 familiar faces from the two networks to unfamiliar control faces. Figure 3 shows that an area of the left fusiform gyrus was more active when viewing unfamiliar faces, whereas the posterior cingulate gyrus/precuneus was more active when viewing familiar faces (p < 0.05, FWE-corrected with threshold-free cluster enhancement within an anatomical mask composed of areas previously shown to be relevant for face perception and memory; see Table 2 for cluster information). While perception for different categories of faces is highly dependent on task demands, our findings are consistent with some previous literature examining recognition of familiar faces (Natu & O’Toole, 2011). The fusiform gyrus has been shown to activate significantly less to famous faces than to strangers in the left hemisphere (Gobbini, Leibenluft, Santiago, & Haxby, 2004), and the posterior cingulate/precuneus area is consistently activated more to personally familiar faces when compared to strangers (Gobbini et al., 2004; Pierce, Haist, Sedaghat, & Courchesne, 2004; Sugiura et al., 2001). There were no univariate differences between responses to faces across the two networks.

**Figure 3.**
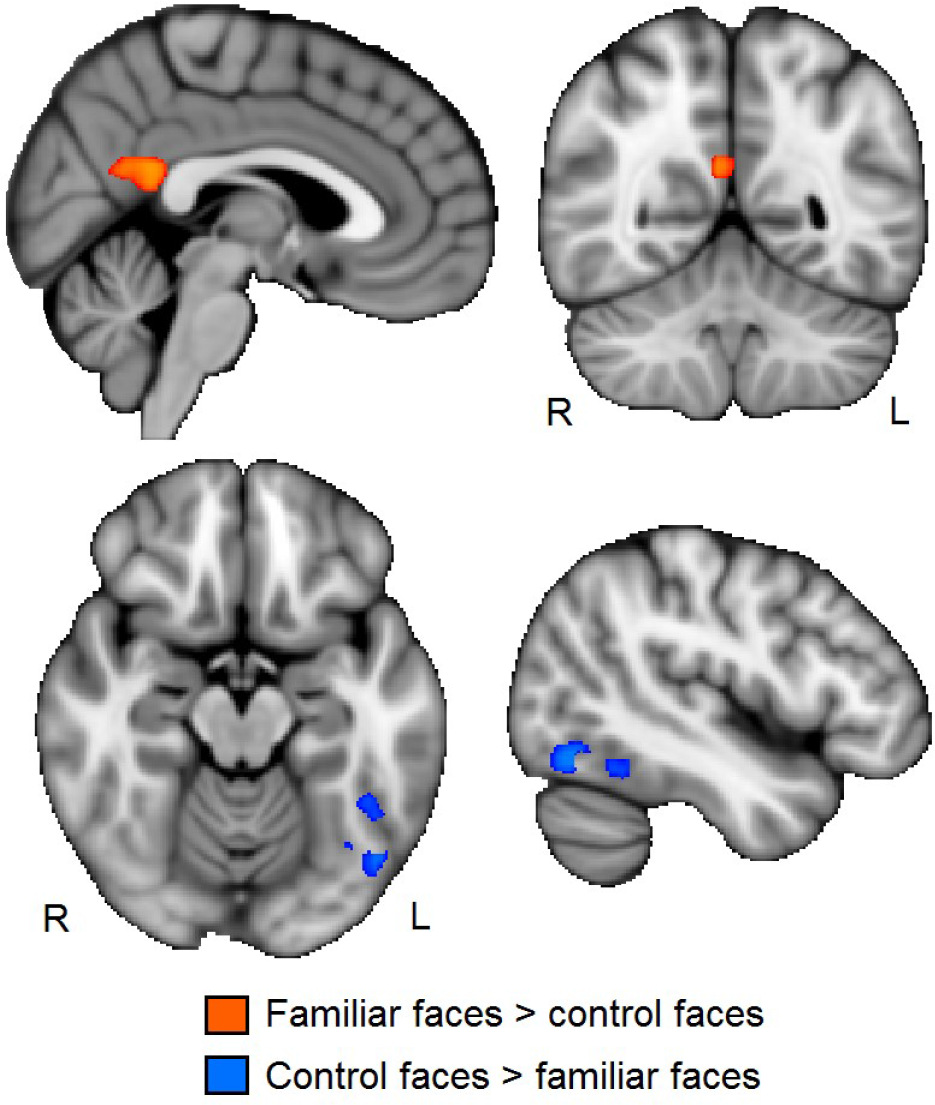
Results from group analysis of z-statistic maps of familiar (network 1 and 2) faces vs. control faces (p < 0.05, FWE-corrected with threshold-free cluster enhancement within an anatomical mask composed of areas previously shown to be relevant for face perception and memory).

**Table 2.** Coordinates, cluster size, and peak activity for the group-level clusters from the univariate familiar vs. control face analysis.

| Cluster | Peak Value (t) | Voxels | x | y | z |
| --- | --- | --- | --- | --- | --- |
| **Familiar > Control** |  |  |  |  |  |
| *Posterior cingulate/precuneus* | 4.44 | 152 | 6 | -48 | 20 |
| *Posterior cingulate/precuneus* | 3.62 | 7 | 2 | -36 | 30 |
| *Posterior cingulate/precuneus* | 3.37 | 6 | -4 | -60 | 32 |
| **Control > Familiar** |  |  |  |  |  |
| *L fusiform gyrus* | 4.01 | 248 | -44 | -68 | -14 |
| *L fusiform gyrus* | 4.11 | 30 | -38 | -90 | -8 |
| *L fusiform gyrus* | 4.19 | 11 | -24 | -30 | -26 |

### Representational Similarity Analysis

To examine whether information related to social network recall is represented in the brain, we carried out RSA searchlight analysis on several DMs representing different types of information about the networks. The first compared neural pattern similarity to social tie strength, with more similar neural responses to any pair of faces representing a closer relationship between those faces. Neural pattern similarity that reflects this network structure would indicate that the brain carries information about the true relationship between individuals, regardless of whether people recall those relationships accurately. We did not find a significant correlation between these measures in our analyses. As the network properties differed between network 1 and 2 (see Methods section for details), we compared the two networks and found no significant differences.

While the pattern similarity to viewing faces was not significantly associated with social tie strength, it was significantly associated with the subjects’ memory for that tie strength. We assessed this by measuring each subject's absolute distance from each true network structure and the 1-correlation distance between the neural response to each face viewed in the scanner. An association between these two measures would indicate that the more accurately a subject perceives the true relationship tie strength between a pair of faces, the more similar their neural pattern response is to those two faces. In other words, this association does not rely on the actual connection strength of the relationships themselves, but the subject's memory of that connection, reflecting a second-order knowledge or understanding of a social relationship. Neural pattern similarity in the left TPJ, the left fusiform gyrus, the subcallosal cingulate cortex, the cerebellum, the left thalamus, and a small portion of the left lateral occipital lobe was significantly correlated with the recall accuracy model, suggesting that neural populations within these areas are important for accurate perception of social relationship strength (Figure 4). Table 3 reports MNI coordinates, cluster size, and peak voxel activity of results. As with tie strength similarity, we compared the two networks to each other separately and found no significant differences. This indicates that the significant findings are not due simply to frequency of the face pairs being presented, as this differed between the two networks.

**Figure 4.**
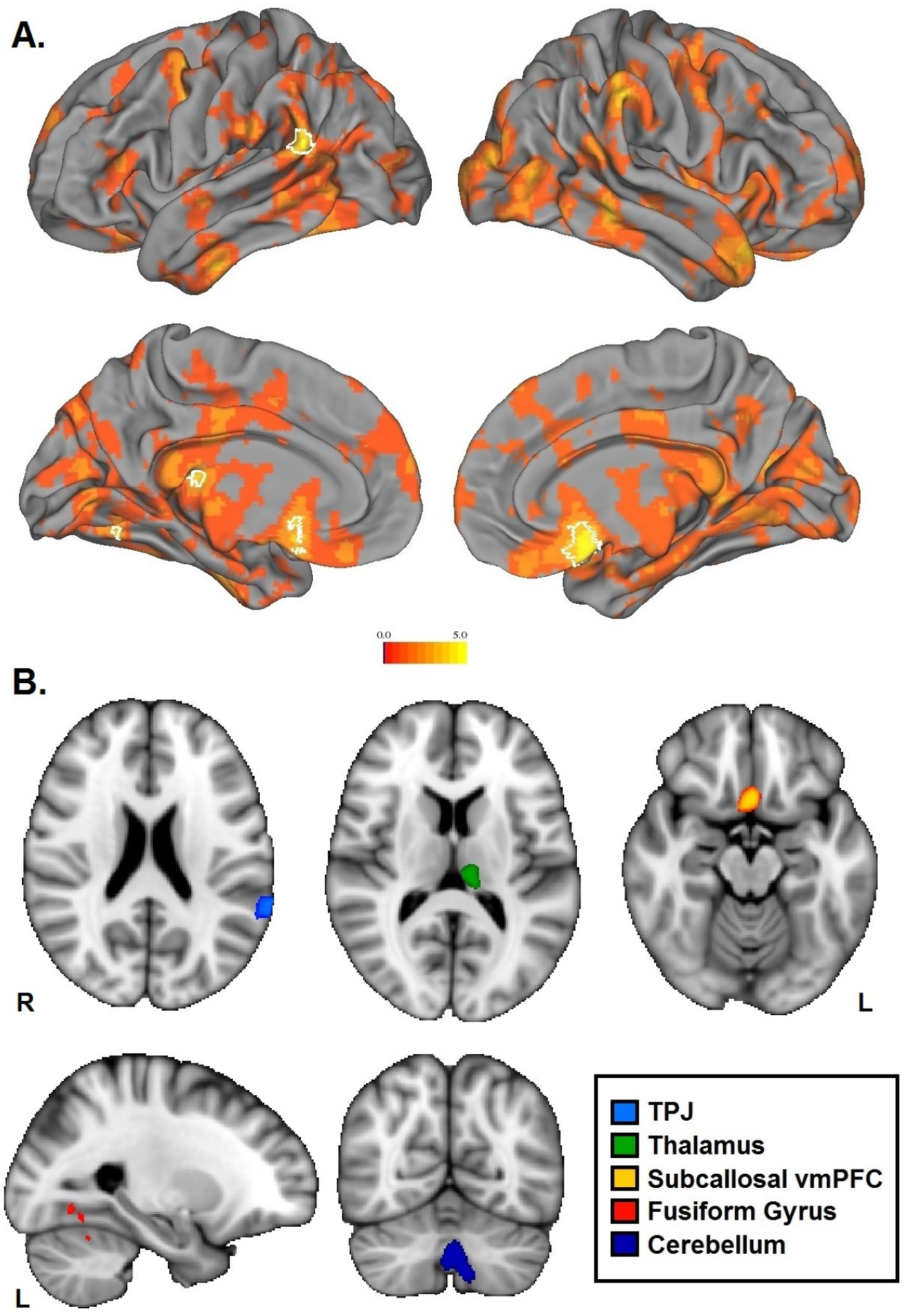
Results from group-level nonparametric 1-sample t-test on the correlation maps from RSA searchlight on the recall accuracy model. A. t-statistic map of all positive t-values projected onto the surface, where white borders delineate significant clusters from the group analysis (p < 0.05, FWE-corrected with threshold-free cluster enhancement). B. The same significant clusters projected in the volume.

**Table 3.** Coordinates, cluster size, and peak activity for the group-level significant clusters from the recall error model.

| Cluster | Peak Value (t) | Voxels | x | y | z |
| --- | --- | --- | --- | --- | --- |
| Cerebellum | 3.92 | 730 | 0 | -64 | -38 |
| Subcallosal vmPFC | 5.6 | 274 | 2 | 14 | -16 |
| Thalamus | 4.14 | 132 | -10 | -28 | 10 |
| TPJ | 4.31 | 117 | -64 | -48 | 18 |
| Fusiform Gyrus | 3.28 | 11 | -26 | -60 | -12 |
| Fusiform Gyrus | 3.5 | 5 | -24 | -66 | -8 |
| Fusiform Gyrus | 3.37 | 5 | -26 | -56 | 26 |
| Lateral Occipital | 3.65 | 3 | -52 | -56 | 2 |
| Lateral Occipital | 3.66 | 3 | -56 | -70 | -2 |

We also conducted RSA searchlights using two other dissimilarity matrix models: recalled structure as measured by behavioral judgments, and the group average of those behavioral judgments (Figure 5). Neural pattern similarity that reflects behavioral recall would indicate that the brain carries information about an individual's perception of relationships, regardless of how accurate those perceptions are. This perceived structure at the group average level can show general trends in how relationships are viewed by groups. However, neither model reached significance in the whole-brain searchlight analysis. Finally, we utilized a separate functional localizer to create regions of interest selective for face processing in the STS and fusiform gyrus, and conducted RSA correlations across each ROI for every subject. No selected regions yielded significant results.

**Figure 5.**
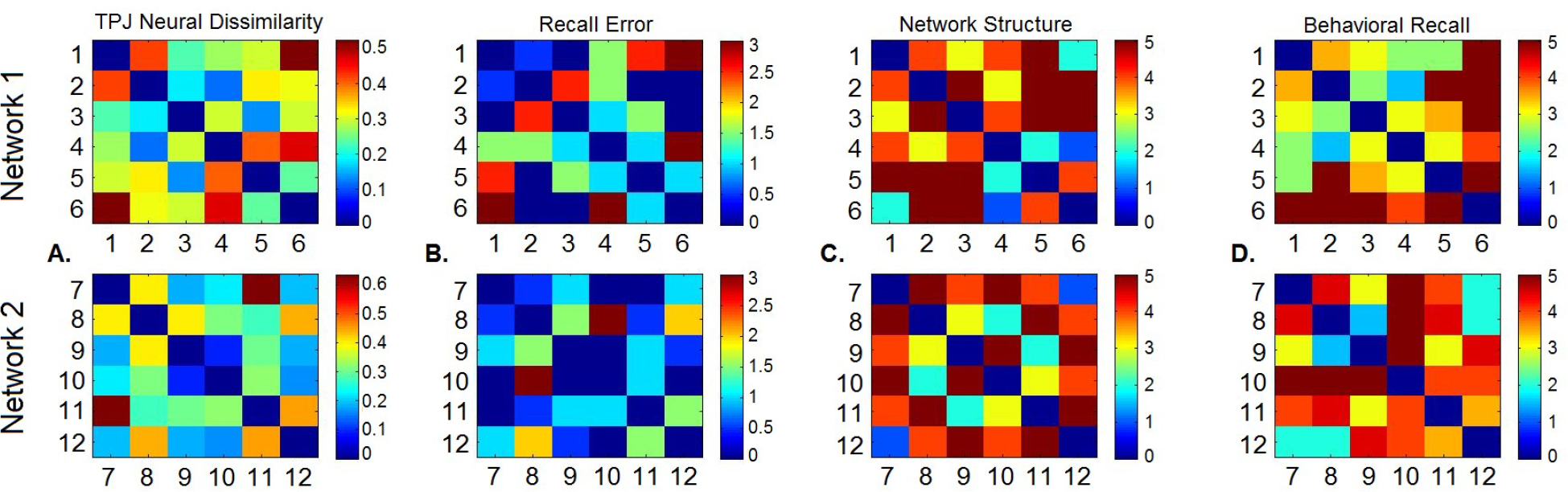
Dissimilarity matrices between face pairs for a sample subject. A. Neural dissimilarity in an example region in the temporoparietal junction. B. Recall error DM (0 = perfectly accurate recall). C. True network structure DM (0 = unconnected). D. Behavioral recall of face pair strength (0 = unconnected).

## Discussion

In this study, we used fMRI and RSA to examine the neural representational space of friendship connections of members of a social network. Indirect connections (i.e., the friends of our friends) play an important role in assessing our own place in our social world. This could include knowledge about social hierarchy which may affect how we act around different network members, or knowledge about which people are more well-connected and might therefore be better to approach for acquiring resources. We examined if the strength of ties between pairs of network members was represented in human brain via the similarity of fMRI responses associated with viewing the faces of those members. We did not find support for this proposal. Instead, our results show that several brain regions, including the TPJ, subcallosal vmPFC, fusiform gyrus, cerebellum, and thalamus, represent memory or knowledge about tie strength, rather than tie strength itself, in the similarity of neural patterns between face pairs. That is, these areas code memory for relationship strength regardless of what that connection is, or even whether there is a connection at all, within a social network. The more accurately a participant recalled the tie strength for a pair of faces (regardless of the closeness of ties), the more similar the pattern of fMRI responses was to viewing those two faces.

Our results indicated that the relationship between memory for tie strength and neural pattern similarity was not due to factors such as the frequency at which different faces were paired with others during the learning of the network, as we found no differences in memory performance or RSA results between a network in which some faces were paired more often with others (variable-centrality network) and one in which all faces had the same number of connections to other network members (fixed-centrality network). In addition, participants saw each individual face the same number of times as they learned one of the two networks and there were no significant behavioral or neural differences between the two networks, and therefore our results are not driven by the familiarity of one face over any other. Instead, our results indicate that information about pairs of network members, such as how close they are, is coded in TPJ and vmPFC via the similarity of neural responses.

Both the TPJ and vmPFC have been consistently linked with complex facets of social understanding, such as tracking the popularity of real-world social network members (Zerubavel, Bearman, Weber, & Ochsner, 2015). The TPJ, dorsomedial PFC, and ventrolateral PFC are engaged when participants recall different facets of socially relevant knowledge (Satpute et al., 2014). The left TPJ is selectively modulated by vasopressin, a neuropeptide linked to a number of complex social behaviors, during social recognition (Zink et al., 2011) and lesions to the left TPJ lead to specific deficits in social reasoning (Samson, Apperly, Chiavarino, & Humphreys, 2004). The vmPFC shows increased activation when thinking about friends compared to kin (Wlodarski & Dunbar, 2016), and the subgenual cingulate cortex is involved in tracking individual differences in perceptions of cohesiveness in kin groups (Rüsch et al., 2014). Our findings are in line with this previous literature showing the importance of these areas in forming and maintaining social relationships. They further indicate that these areas are not only important in the knowledge and understanding of other individuals, but they also support learning relations between individuals in groups. The fusiform gyrus is also heavily involved in social perception, particularly in response to face stimuli (Kanwisher, McDermott, & Chun, 1997). While early models of face perception suggested a strict feed-forward mechanism for distinguishing, identifying, and gaining socially-relevant information from faces, recent proposals indicate a more interactive process between different neural regions when engaging in higher-order social face perception (Atkinson & Adolphs, 2011). Our data indicates that patches of the fusiform gyrus do not simply perceive and distinguish facial features (from each other as well as non-face stimuli), but are also involved in learning more abstract social relationships between faces.

A large meta-analysis of fMRI studies has revealed that areas of the cerebellum are activated in several features of social cognition, with increases in activity occurring with increasing social abstraction levels in the cognitive tasks (Overwalle, Baetens, Marien, & Vandekerckhove, 2014). The authors suggest this cerebellar activity is due to a general increase in cognitive task demands, in line with the theory of the cerebellum as a cognitive process modulator (Andreasen & Pierson, 2008). Our finding that the cerebellum is involved in accurate knowledge of abstract learned relationships between others is consistent with this. Furthermore, we found that the thalamus is also involved in this process. The thalamus has a large number of connections to other areas of the brain, and has been shown to have specific emotional and socially-relevant associations (Christoffel et al., 2015; Feng et al., 2016; Ioannidis et al., 2013). It also has high functional connectivity to the hippocampus (Stein et al., 2000), and may be a critical link in the formation of episodic memories, regardless of the sociality of those memories (Aggleton et al., 2010).

The findings of the present study complement a recent paper by Parkinson and colleagues (2017), who reported that neural pattern similarity in ventral mPFC and lPFC, TPJ and lateral temporal cortex, as well as other regions, to viewing videos of individuals from a participants’ social network was associated with network characteristics of those viewed, including centrality within the network, social distance from the participant, and the ‘brokerage’ of an individual (the extent to which they connected other, low contact network members to others in the network). Parkinson and colleagues took advantage of the one, real-world social network in which all of the participants and those who were used as stimuli were embedded. In contrast, we used an artificial social network in which all network members were initially unfamiliar to the participants, and thus only examined relationships between the network members, and not those between network members and our participants. Together, the more naturalistic, field-work informed approach of Parkinson and colleagues and the laboratory-based approach of as ours, in which factors such as familiarity and the statistics of connections were experimentally controlled, both reveal that social network information is represented in brain regions implicated in social cognition through the similarity of local patterns of neural responses to viewing individual network members.

While most of our subjects were able to accurately report relationship ties, there were individual differences between ability to recall relationship strength (measured by the correlation between the true structure and the reported structure of the networks). Previous literature does indicate that there are individual differences in social recall. Individuals tend to report group and relationship averages or norms more accurately than individual interactions, but more experienced observers show more accurate recall, especially when group structure is transitive (Freeman & Romney, 1987; Freeman, 1992; Kumbasar, Romney, & Batchelder, 1994). It has been suggested that humans use cognitive heuristics such as triadic closure in order to remember social ties (De Soto, 1960; Freeman, 1992; Brashears, 2013; Brashears & Quintaine, 2015). Overestimation of symmetric ties for less central network members, and underestimation of more central network members, has also been reported previously (Krackhardt, 1987). There are also differences in the ability to perceive and remember non-social patterns, but evidence suggests that learning, remembering, and storing social information might be distinct from traditional learning and memory systems (Okuyama, Kitamura, Roy, Itohara, & Tonegawa, 2016; Meyer, Taylor, & Lieberman, 2015; Tendler & Wagner, 2015). Further experiments could explore this type of task explicitly, as prior social network learning studies informed participants that they would be tested on connections (Brashears, 2013; De Soto, 1960).

The way in which people learn and remember social connections between individuals in groups has a considerable impact on everyday life. We are not only able to perceive and understand the social signals of other individuals, but we can also perceive and understand information about social connections or relationships in which we are not directly involved. Our results show that representations of these indirect connections are coded in the pattern of neural responses associated with viewing related individuals. This is a critically important skill because the accuracy with which we perceive and remember subtle connections and relationships seen in our surroundings helps us move more freely and easily in our highly social world.

## Acknowledgments

This research was funded by Office of Naval Research Award N00014-10-1-0198.

